# Ultrastructure of the axonal periodic scaffold reveals a braid-like organization of actin rings

**DOI:** 10.1101/636217

**Authors:** Stéphane Vassilopoulos, Solène Gibaud, Angélique Jimenez, Ghislaine Caillol, Christophe Leterrier

## Abstract

Recent super-resolution microscopy studies have unveiled a periodic scaffold of actin rings regularly spaced by spectrins under the plasma membrane of axons. However, ultrastructural details are unknown, limiting a molecular and mechanistic understanding of these enigmatic structures. Here, we combine platinum-replica electron and optical super-resolution microscopy to investigate the cortical cytoskeleton of axons at the ultrastructural level. Immunogold labeling and correlative super-resolution/electron microscopy allow us to unambiguously resolve actin rings as braids made of two long, intertwined actin filaments connected by a dense mesh of aligned spectrins. This molecular arrangement contrasts with the currently assumed model of actin rings made of short, capped actin filaments. Along the proximal axon, we resolved the presence of phospho-myosin light chain and the scaffold connection with microtubules via ankyrin G. We propose that braided rings explain the observed stability of the actin-spectrin scaffold and ultimately participate in preserving the axon integrity.

## Introduction

Neurons develop an intricate axonal arborization to ensure the proper flow and processing of information. This extraordinary architecture is built and maintained by a unique organization of the axonal cytoskeleton (Leterrier et al., 2017; Tas and Kapitein, 2018). Pioneering electron microscopy (EM) studies have revealed parallel arrays of microtubules within axons that support vesicular transport over long distances (Peters et al., 1991). Only recently has optical super-resolution microscopy been able to unveil striking actin assemblies within axons (Papandréou and Leterrier, 2018; Sigal et al., 2018): the axon is lined by a membrane-associated periodic scaffold (MPS) composed of circumferential actin rings spaced every ∼185 nm by tetramers of spectrins (D’Este et al., 2015; Leterrier et al., 2015; Xu et al., 2013; Zhong et al., 2014). A molecular and mechanistic understanding of actin rings and the MPS is still lacking, as EM has not yet visualized this assembly despite its presence in virtually all neurons (D’Este et al., 2016; Dubey et al., 2018; He et al., 2016; Unsain et al., 2018b). We thus set out to investigate the ultrastructure of the axonal actin-spectrin scaffold, performing EM observation of the MPS and combining it with optical super-resolution microscopy to reveal its molecular architecture.

## Results

### Mechanical unroofing of cultured neurons preserves the MPS

To obtain three-dimensional views of the axonal cortical cytoskeleton in a close to native state, without the need for detergent extraction or sectioning (Jones et al., 2014; Schrod et al., 2018), we mechanically unroofed cultured hippocampal neurons using ultrasound (Heuser, 2000). We first verified that the periodic organization of the MPS was still detected along the axon of unroofed neurons by super-resolution fluorescence microscopy based on single-molecule localization (SMLM). Staining of unroofed cultures without permeabilization highlighted the unroofed neurons with accessible ventral membrane (Fig. 1a). In unroofed proximal axons, we detected ∼180 nm periodic rings for actin (Fig 1b). A similar ∼190 nm periodic pattern was also obtained when labeling the carboxyterminus of ß4-spectrin along the axon initial segment (AIS, Fig. 1c) and ß2-spectrin along the more distal axon, although it was more difficult to unroof distal processes (Fig. 1d). A strong periodic pattern resulted from combining antibodies against axonal spectrins (α2, ß2 and ß4, Fig. 1e) (Galiano et al., 2012). We measured no difference with non-unroofed, permeabilized neurons (Supplementary Figure 1a-g) after quantification of the MPS spacing and periodicity using autocorrelation measurements from SMLM images (Fig. 1f-h), except a more marked periodicity of actin along the AIS as unroofing removed the intra-axonal actin patches (Watanabe et al., 2012).

**Fig. 1.**
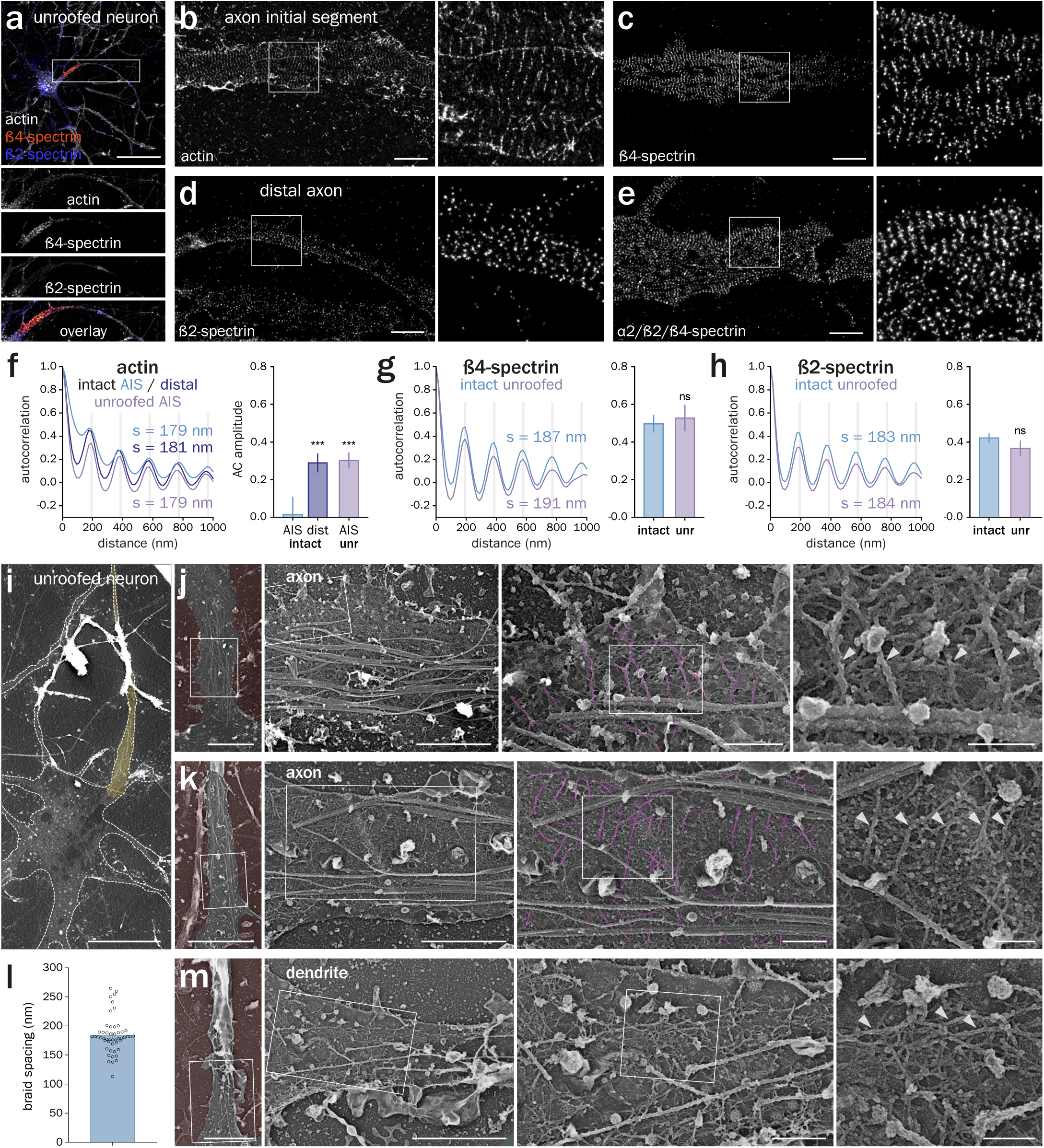
The actin-spectrin MPS is conserved in unroofed axons and visible by PREM. **(a)** Epifluorescence image of an unroofed neuron labeled for actin (gray), ß4-spectrin (orange) and ß2-spectrin (blue). **(b-e)** SMLM images showing the periodic pattern of actin (b), ß4-spectrin (c), ß2-spectrin (d) and α2/ß2/ß4-spectrin (e) along unroofed axons. **(f-h)** Left, autocorrelation curve of the labeling for actin (f), ß4-spectrin (g), or ß2-spectrin (h). Spacing (s) is indicated. Right, measurement of corresponding the autocorrelation amplitude. **(i)** Low-magnification PREM view of an unroofed neuron and its axon (yellow). **(j-k)** PREM views of an unroofed axon showing the regularly spaced braids (magenta, arrowheads) perpendicular to microtubule fascicles. **(l)** Distance between regularly spaced actin braids in axons measured on PREM views. **(m)** PREM view of an unroofed dendrite from the same neuron shown in (k) containing mostly longitudinal actin filaments (arrowheads). Scale bars 40 µm (a), 2 µm (b-e), 20 µm (i), 5, 2, 0.5 and 0.2 µm (j-k & m, left to right).

### Platinum-replica EM resolves actin rings as braids of actin filaments

We then imaged unroofed neurons by transmission EM of metal replicas (Heuser and Kirschner, 1980). To identify the axonal process stemming from unroofed cell bodies, we located the characteristic fascicles of AIS microtubules (yellow, Fig. 1i) (Leterrier, 2018). High-magnification views of the axonal membrane-associated cytoskeleton revealed the presence of long, parallel actin filaments, oriented perpendicular to the axis of the axon and to microtubules, and often regularly spaced by ∼200 nm. These structures resembled braids made of two actin filaments (magenta, Fig 1j-k, Supplementary Figure 1h, Supplementary Movies S1-S3). Their average spacing of 184 ± 4 nm (Fig 1l), and their detection only along the axon (Fig. 1m) strongly suggest that these braids are the periodic actin rings observed by SMLM.

To show that these braids were indeed made of actin filaments, we labeled axonal actin for platinum-replica EM (PREM) using two complementary methods. First, we used phalloidin-Alexa Fluor 488 staining followed by immunogold labeling (Jones et al., 2014) and observed numerous gold beads decorating the actin braids (Fig. 2a). Second, we used myosin subfragment 1 (S1) to decorate actin into a ropelike double helix (Heuser and Kirschner, 1980). Myosin S1 extensively decorated the actin braids (Fig. 2b) and was able to protect them during fixation and processing, as their average observable length went from 0.68 ± 0.04 µm to 1.13 ± 0.04 µm (Fig. 2C).

**Fig. 2.**
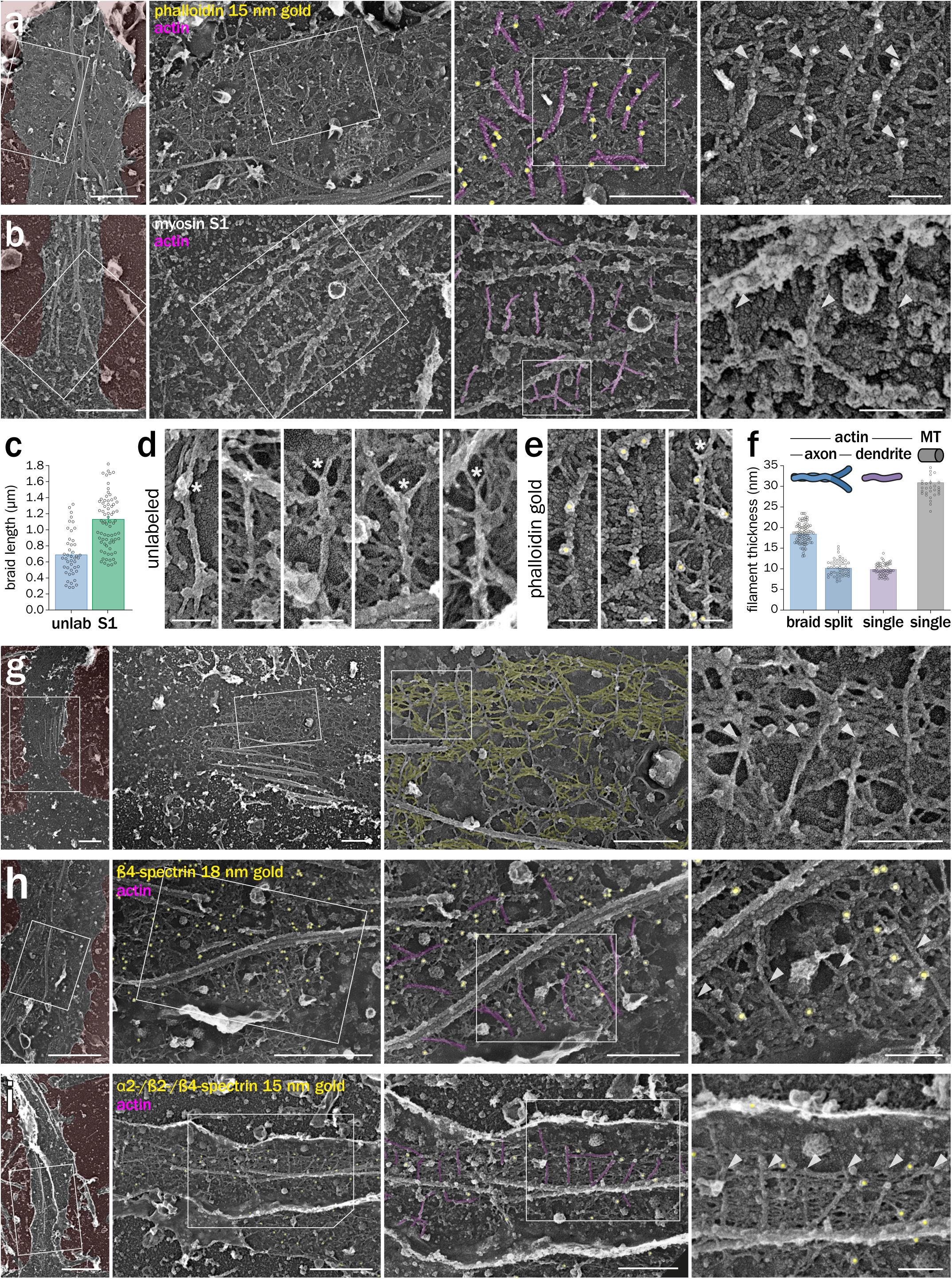
Actin rings are braids of long filaments connected by a spectrin mesh. **(a)** PREM views of axonal actin braids (magenta, arrowheads) labeled with fluorescent phalloidin and immunogold detection of the fluorophore (15 nm gold beads, pseudo-colored yellow). **(b)** PREM views of myosin S1-treated axonal actin braids (magenta, arrowheads). **(c)** Length of the actin braids measured on PREM views in unlabeled (unlab) and myosin S1-labeled (S1) axons. **(d-e)** High-magnification views of individual unlabeled (d) and immunogold-labeled (e) actin braids showing Y bifurcations (asterisks). **(f)** Thickness of filaments measured on PREM views: axonal actin braids before (braid) and after (split) splitting (blue), dendritic single actin filaments (purple) and axonal microtubules (gray). **(g)** PREM views of unroofed axons showing the mesh (yellow) connecting actin braids. **(h)** PREM views of unroofed axons immunogold-labeled (yellow) for ß4-spectrin between actin braids (magenta, arrowheads). **(i)** PREM views of unroofed axons simultaneously immunogold-labeled (yellow) for α2/ ß2/ß4-spectrin between actin braids (magenta, arrowheads). Scale bars 2 µm, 1, 0.5, 0.2 µm (a-b, g-i, left to right), 0.1 µm (d-e).

High-magnification views of individual actin braids confirmed that braids are most often composed of two long (> 0.5 µm) and intertwined actin filaments (Fig. 2d-e). The filaments sometimes separate at the visible extremity of a braid, bifurcating in a Y shape (Fig. 2d-e). This could result from the unwinding of a ring broken into braids during the unroofing procedure, as it is rarer along the myosin S1-protected braids. Alternatively, it may be a snapshot of actin filaments assembling into rings, as ring branching has been described along the proximal axon of living neurons (D’Este et al., 2015). We then confirmed that braids are made of two actin filaments by measuring the diameter of individual actin braids before and after splitting (Fig. 2f): braids are 18.2 ± 0.3 nm thick. When splitting, they form two 10.2 ± 0.3 nm filaments, similar in diameter to single actin filaments in dendrites (9.9 ± 0.2 nm), and to the reported value of 9-11 nm for a single actin filament rotary-replicated with ∼2 nm of platinum (Heuser, 1983; Heuser and Kirschner, 1980). The organization of rings as braids of long, unbranched filaments is to our knowledge unique among cellular actin structures (Blanchoin et al., 2014) and contrasts with the currently assumed view that axonal actin rings made of short, capped and bundled actin filaments (Leite et al., 2016; Xu et al., 2013).

### Actin braids are connected by an organized mesh of spectrins

Between actin rings, the periodic organization of spectrins implies that 190-nm long spectrin tetramers connect two adjacent rings (Supplementary Figure 1f-g) (Bennett et al., 1982; Xu et al., 2013). On our PREM images, actin braids are connected by a dense mesh of rods aligned perpendicular to the braids (yellow, Fig. 2g, Supplementary Movie S4). To identify the components of this mesh, we used immunogold labeling against ß4-spectrin. Gold beads decorated the mesh filaments in-between actin braids (Fig. 2h). The immunolabeling efficiency along the unroofed proximal axons was low for ß2-spectrin, as it is mostly present along the distal axon (Supplementary Figure 2a). Similarly, we could not reliably detect adducin, as it is mostly absent from the AIS and proximal axon, as previously reported (Jones et al., 2014). To better detect spectrins and confirm that they form the connecting mesh, we used a triple immunogold labeling of *α*2-, ß2-and ß4-spectrin. This resulted in a number of gold beads decorating the submembrane mesh, often half-way between the actin braids (Fig. 2i). Overall, the labeling experiments demonstrate that PREM resolves axonal actin rings made from pairs of long, intertwined filaments, and reveal the dense alignment of spectrins that connect these rings to form the axonal MPS.

### Ultrastructural organization of proximal MPS components

We next took advantage of PREM of unroofed neurons to determine the ultrastructural localization of MPS components known to specifically localize along the AIS (Leterrier, 2018). The phosphorylated myosin light-chain (pMLC), an activator of myosin II, was recently reported to associate with the MPS along the initial segment (Berger et al., 2018), together with its partner tropomyosin (Abouelezz et al., 2019b). We found concentrated pMLC at the AIS, with areas showing a ∼200 nm periodic pattern on SMLM images (Fig. 3a-b). After unroofing, a fraction of the pMLC staining was retained, although a periodic pattern was not easily detectable (Fig. 3c-d). Immunogold labeling for pMLC followed by PREM showed gold beads often appearing along filaments perpendicular to actin braids (Fig. 3e, brackets). These filaments could be myosins associated with the MPS, and suggest that myosins can crosslink neighboring rings (Wang et al., 2018).

**Fig. 3.**
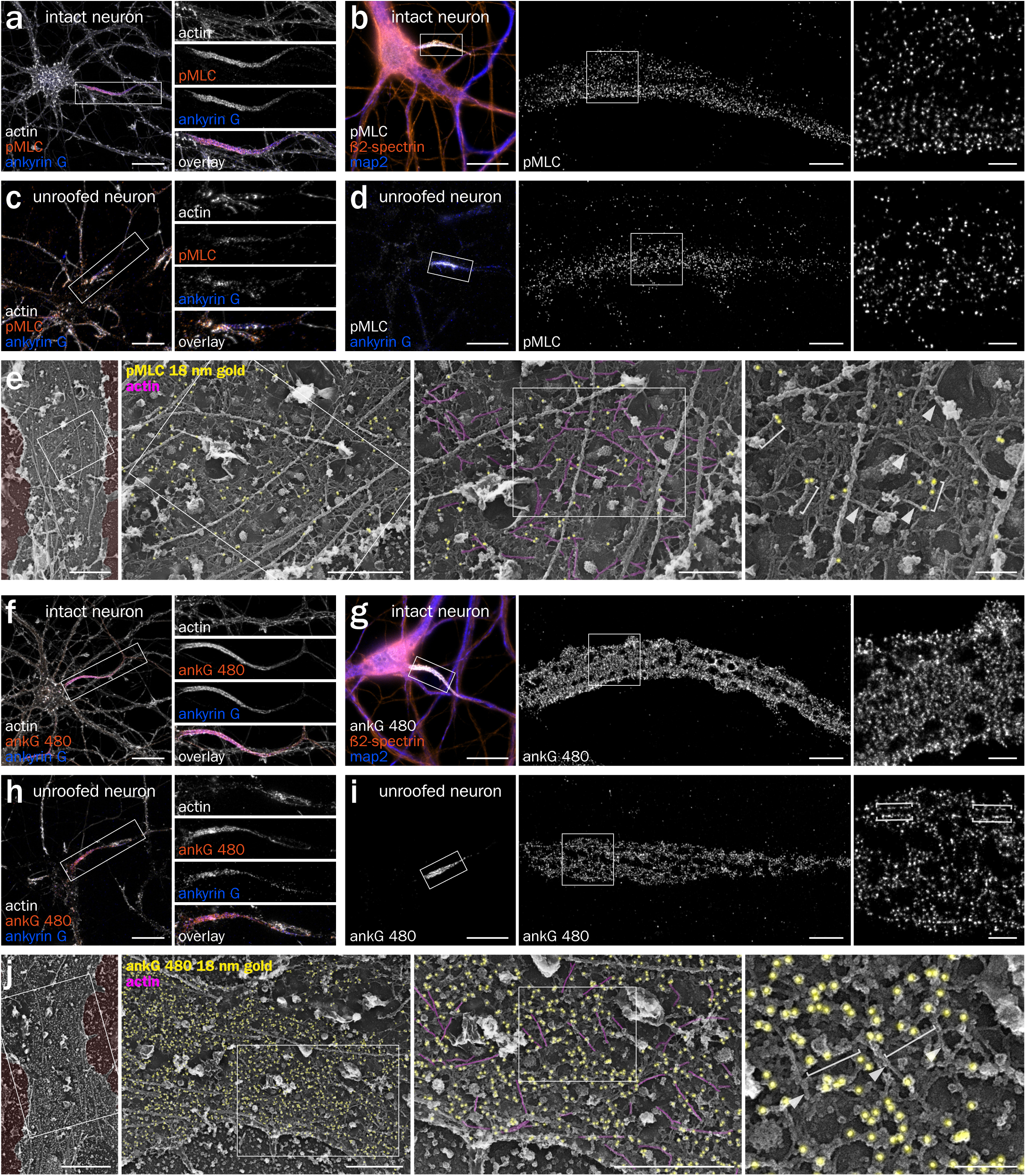
Localization of pMLC and ankyrin G at the AIS of unroofed neurons. **(a)** Epifluorescence image of an intact neuron labeled for actin (gray), pMLC (orange) and ankyrin G (blue). **(b)** SMLM images showing the pattern of pMLC along an intact AIS. **(c)** Epifluorescence image of an unroofed neuron labeled for actin (gray), pMLC (orange) and ankyrin G (blue). **(d)** SMLM images showing the pattern of pMLC along an unroofed AIS. **(e)** PREM views of an unroofed AIS immunogold-labeled (yellow) for pMLC (brackets) apposed to actin braids (magenta, arrowheads). **(f)** Epifluorescence image of an intact neuron labeled for actin (gray), 480 kDa ankyrin G tail (ankG 480, orange) and ankyrin G (blue). **(g)** SMLM images showing the pattern of ankG 480 along an intact AIS. **(h)** Epifluorescence image of an unroofed neuron labeled for actin (gray), ankG 480 (orange) and ankyrin G (blue). **(i)** SMLM images showing the pattern of ankG 480 along an unroofed AIS, delineating the profiles of putative microtubules (brackets). **(j)** PREM views of an unroofed AIS immunogold-labeled (yellow) for ankG 480 appearing along spectrin filaments (brackets) in between actin braids (magenta, arrowheads). Scale bars 20 µm (a, c, f, h); 2 µm, 0.5 µm (b, d, g, i, left to right); 2 µm, 1, 0.5, 0.2 µm (e, j, left to right).

Next, we examined the localization of ankyrin G (ankG), a large scaffold protein that recruits other components during AIS assembly (Hedstrom et al., 2007; Leterrier, 2018). Immunolabeling in non-permeabilized conditions after unroofing could not detect the aminoterminal part of ankG or associated AIS membrane proteins such as Nav channels. We thus focused on the carboxyterminal part of ankG that extends below the MPS (Leterrier et al., 2015) and interacts with microtubule-associated End-Binding proteins (EBs) (Leterrier et al., 2011) via SxIP motifs in its tail domain (Fréal et al., 2016). Labeling of the 480 kDa ankG tail (Jenkins et al., 2015) resulted in dense staining along the AIS in intact and unroofed neurons (Fig. 3f-i), often showing a pattern of longitudinal “tracks” along unroofed axons (Fig. 3i, brackets). Immunogold labeling and PREM showed a high concentration along the AIS, with beads densely decorating the spectrin mesh (Fig. 3j). AnkG labeling could be detected along the connecting spectrins in-between actin braids (Fig 3j), consistently with the current model of AIS architecture (Leterrier et al., 2015). Overall, these experiments show how PREM on unroofed neurons provides an unprecedented access to the AIS architecture and can visualize identified proteins within its membrane-bound scaffold.

### Impact of actin perturbations on the ultrastructure of the MPS

We next assessed the effect of drugs that potently target actin assemblies on the MPS organization down to the ultrastructural level. Due to the reported stability of the actin rings (Abouelezz et al., 2019a; Leterrier et al., 2015), we used swinholide A, a drug that inhibits actin polymerization and severs existing filaments (Spector et al., 1999). We also used cucurbitacin E, a drug that inhibits filament depolymerization, stabilizes them, and is compatible with phalloidin staining (Sörensen et al., 2012). At the diffraction-limited level, acute swinholide treatment resulted in a near disappearance, while cucurbitacin induced a marked increase, of the actin staining throughout neuronal processes, with a moderate effect on spectrins labeling (Supplementary Figure 3a-b). Using SMLM, we found that swinholide could only partially disorganize the MPS along the AIS, with a periodic appearance of the remaining actin and of ß4-spectrin (Fig. 4a-b, Supplementary Figure 3c-d), whereas in the distal axon both actin and ß2-spectrin lost their periodicity (Fig. 4c-d, Supplementary Figure 3e-f). This higher stability of the MPS in the initial segment compared to the distal axon is consistent with previous results (Abouelezz et al., 2019a; Leterrier et al., 2015) and may be caused by the high density of ankG/membrane protein complexes anchored within the initial segment MPS (see Fig. 3j).

**Fig. 4.**
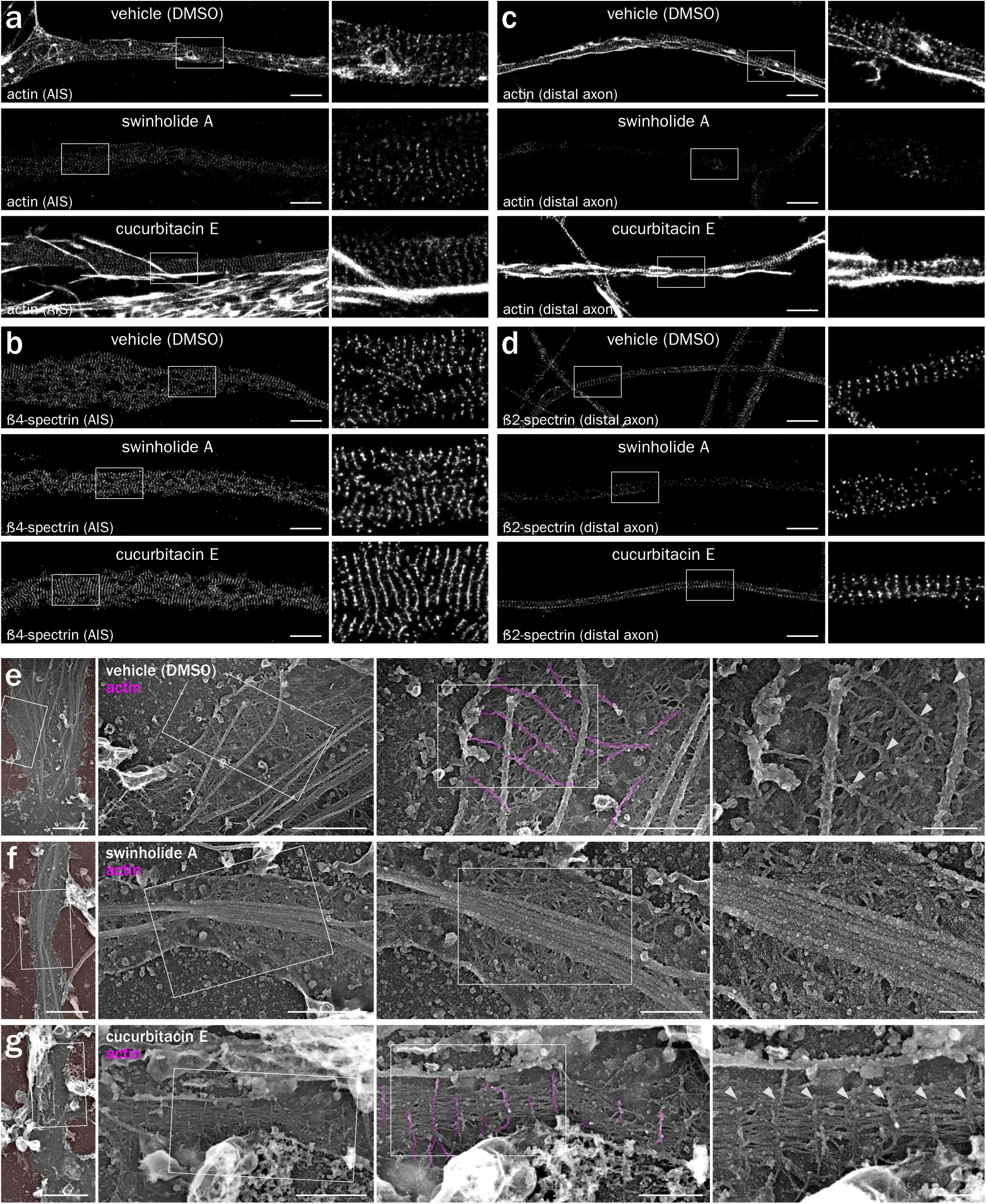
Actin perturbation impacts the MPS ultrastructure. **(a-d)** SMLM images of AIS and distal axons treated for 3h with vehicle (DMSO 0.1%), swinholide A (100 nM) or cucurbitacin E (5 nM), labeled for actin (a and c), ß4-spectrin (b) and ß2-spectrin (d). **(e-g)** PREM views of unroofed axons from neurons treated with vehicle (e), swinholide A (f) or cucurbitacine E (g) showing the presence or absence of actin braids (magenta, arrowheads). Scale bars 2 µm (a-d), 2 µm, 1, 0.5, 0.2 µm (e-g, left to right).

Stabilization by cucurbitacin resulted in the presence of numerous bright longitudinal actin bundles and clusters along the axon, making quantification of the eventual ring stabilization difficult (Fig. 4a and 4c, Supplementary Figure 3c and 3e). Cucurbitacin did not change the ß4-spectrin pattern, but ß2-spectrin showed more consistent bands and a larger autocorrelation amplitude of the pattern (Fig. 4b & 4c, Supplementary Figure 3d and 3f) (Unsain et al., 2018a). PREM views of treated, unroofed neurons (Fig. 4e) showed that swinholide induced a disappearance of the transverse actin braids, while the spectrin mesh remained visible along the proximal axon but was partially disorganized, consistent with the SMLM results (Fig. 4f). The reinforcement of the MPS induced by cucurbitacin was visible by PREM, with the occurrence of very regular stretches of actin braids connected by a densified mesh of spectrins (Fig. 4g). In addition to the similar effect detected by SMLM and PREM, these experiments show that the stable actin braids/rings can be modulated by acute perturbations, suggesting a regulated assembly and turnover.

### Resolving actin rings by correlative SMLM/PREM

The braids seen by PREM are not as long and continuous along the axon as the actin rings visible in SMLM (compare Fig. 1b and 1j-k). To explain this, we performed correlative SMLM and PREM on unroofed neurons. After sonication, fixation and labeling for actin and spectrin, proximal axons were imaged by SMLM before being replicated, relocated on the EM grid and observed (Sochacki et al., 2014). The nanoscale precision provided by both techniques allows to closely register the SMLM images and PREM views. Phalloidin staining of actin rings on SMLM images (Fig. 5a and 5d) frequently corresponded with the braids visible on PREM views (Fig. 5b and 5e). A few actin filaments on PREM images were only faintly stained by phalloidin, likely due to the fast labeling protocol used for our correlative approach. SMLM often showed single rings across the whole width of the axon, whereas they only appeared intermittently on the PREM views (Fig. 5c and 5f, Supplementary Movie 5). This may result from ultrastructural damage during SMLM imaging or sample processing for subsequent PREM. However, close examination of PREM views -that only visualize the surface of the replicated sample -suggests that phalloidin-stained actin rings are often embedded between the plasma membrane and the spectrin mesh, and thus partially hidden from PREM view (Fig.5c and 5f). Phalloidin staining also revealed that rings were continuous under microtubules, although they are similarly hidden from PREM view (Fig. 5c and 5f).

**Fig. 5.**
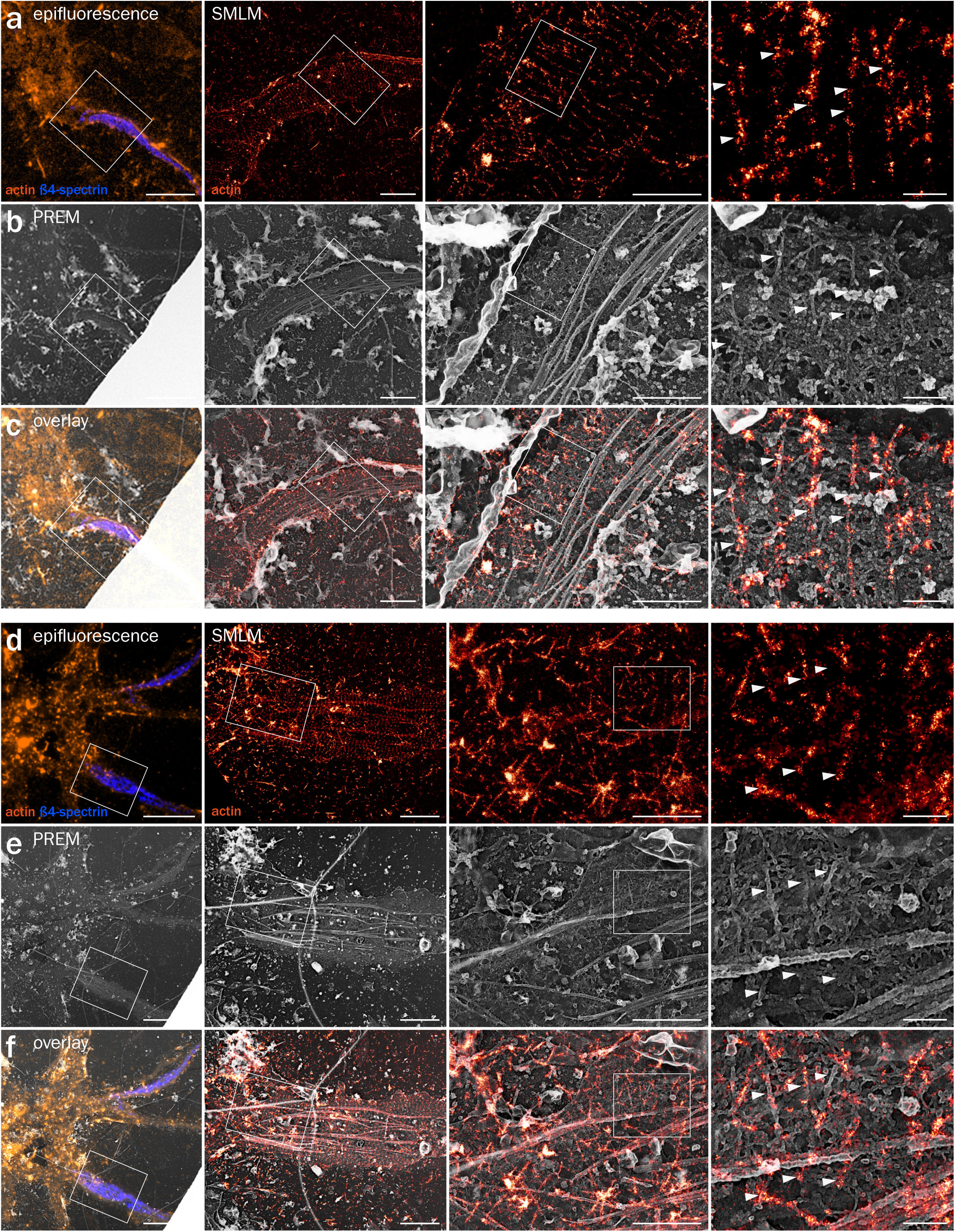
Correlative SMLM/PREM resolves the ultrastructure of actin rings. **(a)** Left, epifluorescence image of an unroofed neuron labeled for actin (orange) and ß4-spectrin (blue). Right, SMLM images of the unroofed proximal axon labeled for actin. **(b)** Corresponding PREM views of the same unroofed neuron and axon. **(c)** Overlay of the SMLM image and PREM views showing the correspondence between actin rings in SMLM and braids in PREM (arrowheads). **(d-f)** Correlative SMLM/PREM images similar to (a-c) for an additional unroofed axon. Scale bars 20, 2, 1, 0.2 µm (from left to right).

### Identification of MPS components by correlative SMLM/PREM

We applied the same strategy to localize spectrins and other MPS components along unroofed axons. SMLM of unroofed neurons labeled for ß4-spectrin using an antibody labeling the center of the spectrin tetramer (Leterrier et al., 2015) exhibited well-defined bands along the AIS (Fig. 6a, Supplementary Figure 4a and 4d). These bands corresponded to the spectrin mesh connecting actin braids on PREM views (Fig. 6b, Supplementary Figure 4b and 4e), and often localized in-between actin braids (Fig. 6c, Supplementary Figure 4b and 4e, Supplementary Movie 6). When imaging ß2-spectrin, the labeling along the proximal axon was fainter by SMLM (Supplementary Fig. 5a), consistent with the presence of ß2-spectrin along the more distal axon (Galiano et al., 2012). This labeling localized to the spectrin mesh in the corresponding PREM views (Supplementary Figure 5b-c, Supplementary Movie 7). We also located pMLC by correlative SMLM/PREM and could resolve spots of pMLC labeling by SMLM (Supplementary Figure 6a). Interestingly, these spots were often present as pairs, associated with thicker filaments roughly perpendicular to actin braids on PREM views (Supplementary Figure 6b-c, brackets), in line with the putative myosin labeling obtained by immunogold labeling (see Fig. 3e, brackets).

**Fig. 6.**
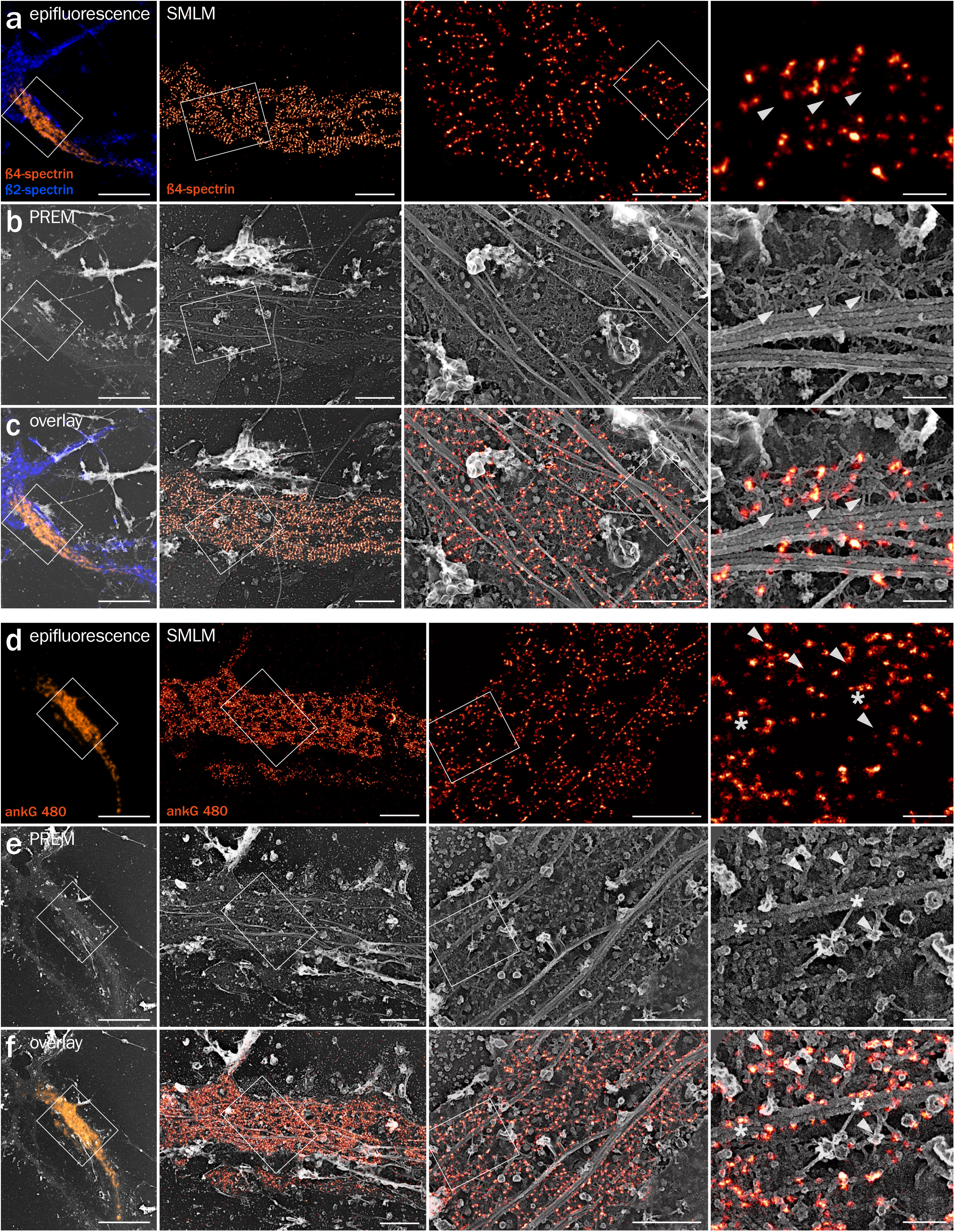
Correlative SMLM/PREM of ß4-spectrin and ankyrin G. **(a)** Left, epifluorescence image of an unroofed neuron labeled for ß4-spectrin (orange) and ß2-spectrin (blue). Right, SMLM images of the unroofed proximal axon labeled for ß4-spectrin. **(b)** Corresponding PREM views of the same unroofed neuron and axon. **(c)** Overlay of the SMLM image and PREM views showing the intercalation between spectrin tetramer centers in SMLM and actin breads in PREM (arrowheads). **(d-f)** Correlative imaging of 480 kDa ankyrin G tail (ankG 480) along an unroofed proximal axon by SMLM and PREM. Actin braids are indicated by arrowheads, and contacts between ankyrin G and microtubules are shown by asterisks. Scale bars 20, 2, 1, 0.2 µm (from left to right).

Finally, we correlated labeling for the 480 kDa ankG tail with PREM views of the same proximal axon (Fig. 6d-f). The density of the labeling resulted in a mesh largely covered by primary and secondary IgGs (Fig. 6e). Correlative overlay revealed that the longitudinal “tracks” (empty lines flanked by a high density of ankG labeling, see Fig. 3i) that we observed on SMLM images correspond to microtubules decorated on both sides with fluorescent molecules that likely correspond to ankG tails (Fig. 6f). This is the first direct visualization of the MPS association with microtubules via ankG along the AIS (Leterrier et al., 2011). This connection could also explain why we usually detected well-preserved actin braids along unroofed areas where micro-tubule bundles are still present (see Fig. 1j-k).

## Discussion

In this work, we obtained the first EM images of the actin-spectrin MPS along axons and provide ultrastructural insight into its molecular organization. Previous work using PREM to visualize the AIS organization did not detect regularly spaced actin filaments or other ultrastructural signs of the MPS (Jones et al., 2014). However, this work used live extraction by a detergent before fixation, an approach that was later shown to disrupt actin rings and the MPS (Zhong et al., 2014). The fact that mechanical unroofing preserves the submembrane assemblies better than detergent extraction suggests that their interaction with the membrane is essential for their stability.

We show that axonal actin rings are made of a small number of long, intertwined filaments rather than by a large number of short, bundled filaments as previously hypothesized (Leite and Sousa, 2016; Unsain et al., 2018b; Xu et al., 2013; Zhang et al., 2017). This short filament model was inferred from the presence of adducin at actin rings, at least along the distal axon (Leite et al., 2016; Xu et al., 2013), because adducin was described as a capping protein that can bind to the barbed end of filaments (Kuhlman et al., 1996). However, adducin is also able to associate to the side of filaments, enhancing the lateral binding of spectrin to actin (Gardner and Bennett, 1987). Although we were not able to directly visualize adducin at the ultrastructural level, our results suggest that this lateral binding role is dominant to enhance actin rings interaction with spectrins. Further studies are needed to clarify the precise localization and role of adducin as well as to confirm the localization of myosin filaments associated with the MPS, in order to explain how actin-associated proteins can regulate radial and longitudinal tension along the axon as well as axonal diameter (Leite et al., 2016; Wang et al., 2018).

Finally, the arrangement of actin rings as a braid of long filaments is relevant to the observed stability of the MPS: closely apposed, intertwined filaments are likely to be resistant to severing factors such as ADF/cofilin (Michelot et al., 2007), and further stabilized by their embedding in the spectrin mesh. Rings made of long intertwined filaments are likely to be stiffer than bundled short filaments, hence could form the rigid part of a flexible and resistant scaffold when linked by elastic spectrins (Zhang et al., 2017), explaining the role of the MPS in the mechanical resistance of axons (Krieg et al., 2017). Beyond shedding light on the molecular underlaying of the MPS function, the ultrastructural insights obtained here could help understand its potential dysfunctions in neurodegenerative diseases, where the axonal integrity is often affected first (Encalada and Goldstein, 2014; Sleigh et al., 2019).

## Methods

### Antibodies and reagents

Rabbit polyclonal anti ßIV-spectrin antibody (against residues 2237-2256 of human ßIV-spectrin, 1:800 dilution for immunofluorescence IF, 1:20 for immunogold IG) was a gift from Matthew Rasband (Baylor College of Medicine, Austin, TX). Mouse monoclonal anti ßII-spectrin (against residues 2101-2189 of human ßII-spectrin, 1:100 for IF, 1:20 for IG) was from BD Biosciences (#612563) (Xu et al., 2013). Chicken anti-map2 antibody was from abcam (#ab5392, 1:1000 for IF). Mouse monoclonal antibodies anti ankyrin G (clone 106/65 and 106/36, 1:300 for IF) were from NeuroMab (Leterrier et al., 2015). Rabbit polyclonal anti 480-kDa ankG (residues 2735-2935 of rat 480-kDa ankG, 1:300 for IF, 1:100 for IG) was a gift from Vann Bennett (Duke University, Durham, NC) (Jenkins et al., 2015). Rabbit anti phospho-Myosin Light Chain 2 Thr18/Ser19 (pMLC) was from Cell Signaling Technologies (#3674, 1:50 for IF, 1:20 for IG) (Berger et al., 2018).

Rabbit polyclonal anti-Alexa Fluor 488 antibody was from Thermo Fisher (A11094, 1:20 for IG). Donkey and goat anti-rabbit and anti-mouse secondary antibodies conjugated to Alexa Fluor 488, 555 and 647 were from Life Technologies or Jackson ImmunoResearch (1:200-1:400 for IF). Donkey anti-rabbit and anti-mouse secondary antibodies conjugated to DNA PAINT handles P1 (ATACATCTA) and P3 (TCTTCATTA), respectively, were prepared according to published procedures (Schnitzbauer et al., 2017). Goat anti-rabbit and anti-mouse secondary antibodies conjugated to 15 nm gold nanobeads were from Aurion (#815011 and #815022 respectively, 1:20 for IG) and the ones conjugated to 18 nm gold nanobeads were from Jackson ImmunoResearch (#111-215-144 and #115-215-146 respectively, 1:15 for IG).

Alexa-Fluor 488 and Alexa-Fluor 647 conjugated phalloidins were from Thermo Fisher (#A12379 and #A2287 respectively), Atto488 conjugated phalloidin (#AD488-81) was from At-to-Tec. DMSO (#D2650), swinholide A (#S9810), cucurbitacine E (#PHL800-13), glutaraldehyde (#G5882) were from Sigma. Paraformaldehyde (PFA, #15714, 32% in water) was from Electron Microscopy Sciences. Myosin S1 (from rabbit skeletal fast muscle) was from Hypermol (#9310-01).

### Animals and neuronal cultures

The use of Wistar rats followed the guidelines established by the European Animal Care and Use Committee (86/609/CEE) and was approved by the local ethics committee (agreement D13-055-8). Rat hippocampal neurons were cultured following the Banker method, above a feeder glia layer (Kaech and Banker, 2006). Rapidly, 12 or 18 mm-diameter round, #1.5H coverslips were affixed with paraffine dots as spacers, then treated with poly-L-lysine. Hippocampi from E18 rat pups were dissected, and homogenized by trypsin treatment followed by mechanical trituration and seeded on the coverslips at a density of 4,000-8,000 cells/cm2 for 3 hours in serum-containing plating medium. Coverslips were then transferred, cells down, to petri dishes containing confluent glia cultures conditioned in B27-supplemented Neurobasal medium and cultured in these dishes for up to 4 weeks. For this work, neurons were fixed at 13 to 17 days in vitro, a stage where they exhibit a membrane-associated periodic scaffold (MPS) along virtually all axons (Xu et al., 2013).

### Cell treatments and fluorescence immunocytochemistry

Treatments were applied on neurons in their original culture medium for 3h at 37°C, 5% CO2: DMSO 0.1% (from pure DMSO), swinholide A 100 nM (from 100 µM stock in DMSO), cucurbitacin E 5 nM (from 5 µM stock in DMSO). Stock solutions were stored at −20°C.

Fluorescent immunocytochemistry for epifluorescence microscopy and SMLM was performed as in published protocols (Jimenez et al., 2019). Cells were fixed using 4% PFA in PEM buffer (80 mM PIPES pH 6.8, 5 mM EGTA, 2 mM MgCl2) for 10 minutes at room temperature (RT). After rinses in 0.1M phosphate buffer (PB), neurons were blocked for 2-3h at RT in immunocytochemistry buffer (ICC: 0.22% gelatin, 0.1% Triton X-100 in PB), and incubated with primary antibodies diluted in ICC overnight at 4°C. After rinses in ICC, neurons were incubated with secondary antibodies diluted in ICC for 1h at RT, rinsed and incubated in fluorescent phalloidin at 0.5 µM for either 1h at RT or overnight at 4°C. Stained coverslips were kept in PB + 0.02% sodium azide at 4°C before SMLM imaging. For epifluorescence imaging, coverslips were mounted in ProLong Glass (Thermo Fisher #P36980).

### Unroofing and PREM immunocytochemistry

Unroofing was performed by sonication as previously described (Heuser, 2000). Coverslips where quickly rinsed three times in Ringer+Ca (155 mM NaCl, 3 mM KCl, 3 mM NaH2PO4, 5 mM HEPES, 10 mM glucose, 2 mM CaCl2, 1 mM MgCl2, pH 7.2), then immersed ten seconds in Ringer-Ca (155 mM NaCl, 3 mM KCl, 3 mM NaH2PO4, 5 mM HEPES, 10 mM glucose, 3 mM EGTA, 5 mM MgCl2, pH 7.2) containing 0.5 mg/mL poly-L-lysine, then quickly rinsed in Ringer-Ca then unroofed by scanning the coverslip with rapid (2-5s) sonicator pulses at the lowest deliverable power in KHMgE buffer (70 mM KCl, 30 mM HEPES, 5 mM MgCl2, 3 mM EGTA, pH 7.2).

Unroofed cells were immediately fixed using fixative in KHMgE: 4% PFA for 10 minutes for epifluorescence or SMLM of fluorescence-labeled samples, 4% PFA for 45 minutes for PREM of immunogold-labeled samples, 3% PFA-1% glutaraldehyde or 2% PFA-2% glutaraldehyde for 10 to 20 minutes for PREM and correlative SMLM/PREM. Glutaraldehyde-fixed samples were subsequently quenched using 0.1% NaBH4 in KHMgE for 10 minutes. Immunofluorescence labeling was performed as above, but replacing the ICC buffer with a detergent-free buffer (KHMgE, 1% BSA). Immunogold labeling was performed in the same detergent-free solution: samples were blocked for 30’, incubated 1h30 with the primary antibodies diluted to 1:20, rinsed, incubated two times 20 minutes with the gold-coupled secondary antibodies, then rinsed.

### Actin immunogold and myosin S1 labeling

For immunogold labeling of actin (Jones et al., 2014), unroofed neurons were fixed with 2% PFA in KHMgE, then quenched for 10 minutes in KHMgE, 100 mM glycine, 100 mM NH4Cl. After blocking in KHMgE, 1% BSA, they were incubated with phalloidin-Alexa Fluor 488 (0.5 µM) for 45 minutes, then immunolabeled using an anti-Alexa Fluor 488 primary antibody and a gold-coupled goat anti-rabbit secondary antibody as described above for immunogold labeling.

For myosin S1 labeling, neurons were unroofed in PEM100 buffer (100 mM PIPES-KOH pH 6.9, 1 mM MgCl2, 1 mM EGTA), treated with 0.25 mg/mL myosin S1 in PEM100 buffer for 30 minutes, then fixed with 2% glutaraldehyde in PEM100 buffer for 10 minutes.

### Platinum-replica sample processing

Fixed samples were stored and shipped in KHMgE, 2% glutaraldehyde. Cells were further sequentially treated with 0.5% OsO4, 1% tannic acid and 1% uranyl acetate prior to graded ethanol dehydration and hexamethyldisilazane substitution (HMDS) (Sigma). Dried samples were then rotary-shadowed with 2 nm of platinum and 5-8 nm of carbon using an ACE600 high vacuum metal coater (Leica Microsystems). The resultant platinum replica was floated off the glass with hydrofluoric acid (5%), washed several times on distilled water, and picked up on 200 mesh formvar/carbon-coated EM grids.

### Epifluorescence and SMLM

Diffraction-limited images were obtained using an Axio-Observer upright microscope (Zeiss) equipped with a 40X NA 1.4 or 63X NA 1.46 objective and an Orca-Flash4.0 camera (Hamamatsu). Appropriate hard-coated filters and dichroic mirrors were used for each fluorophore. Quantifications were performed on raw, unprocessed images. An Apotome optical sectioning module (Zeiss) and post-acquisition deconvolution (Zen software, Zeiss) were used to acquire and process images used for illustration.

For single color SMLM, we used STochastic Optical Microscopy (STORM). STORM was performed on an N-STORM microscope (Nikon Instruments). Coverslips were mounted in a silicone chamber filled with STORM buffer (Smart Buffer Kit, Abbelight). The N-STORM system uses an Agilent MLC-400B laser launch with 405 nm (50 mW maximum fiber output power), 488 nm (80 mW), 561 mW (80 mW) and 647 nm (125 mW) solid-state lasers, a 100X NA 1.49 objective and an Ixon DU-897 camera (Andor). After locating a suitable neuron using low-intensity illumination, a TIRF image was acquired, followed by a STORM acquisition. 30,000-60,000 images (256×256 pixels, 15 ms exposure time) were acquired at full 647 nm laser power. Reactivation of fluorophores was performed during acquisition by increasing illumination with the 405 nm laser. When imaging actin, 30 nM phalloidin-Alexa Fluor 647 was added to the STORM buffer to mitigate actin unbinding during imaging (Jimenez et al., 2019).

For two-color SMLM (Fig. 1D-E), we used STORM in combination with DNA-PAINT (Schnitzbauer et al., 2017). Neurons were labeled using secondary antibodies coupled to a PAINT DNA handle for ß4- or ß2-spectrin (rabbit P3 or mouse P1, respectively), as well as phalloidin-Alexa Fluor 647 for actin. Imaging was done sequentially (Jimenez et al., 2019), first for actin in STORM buffer (60,000 frames at 67 Hz), then for ß4- or ß2-spectrin in PAINT buffer (0.1M phosphate buffer saline, 500 mM NaCl, 5% dextran sulfate, pH 7.2) supplemented with 0.12-0.25 nM of the corresponding PAINT imager strand coupled to Atto650 (Metabion, 40,000 frames at 33 Hz). No chromatic aberration correction was necessary as both channels were acquired using the same spectral channel, and translational shift was corrected by autocorrelation and manual refinement.

The N-STORM software (Nikon Instruments) was used for the localization of single fluorophore activations. After filtering out localizations to reject too low and too high photon counts, the list of localizations was exported as a text file. Image reconstructions were performed using the ThunderSTORM ImageJ plugin (Ovesny et al., 2014) in Fiji software (Schindelin et al., 2012). Custom scripts and macros were used to translate localization files from N-STORM to ThunderSTORM formats, as well as automate images reconstruction for whole images and detailed zooms. STORM processing scripts used in this work can be found at https://github.com/cleterrier/ChriSTORM.

### Fluorescence image analysis

Quantification of the labeling intensities on epifluorescence images was done by first tracing the region of interest along a process (AIS, axon or dendrite) using the NeuronJ plugin (Meijering et al., 2004) in Fiji software. Tracings were then translated into ImageJ ROIs and the background-corrected intensities were measured for each labeled channel. Intensity measurement scripts are available at https://github.com/cleterrier/Measure_ROIs.

Quantification of the scaffold periodicity on SMLM images was performed using autocorrelation analysis (Zhong et al., 2014): if an intensity profile is periodically patterned, its autocorrelation curve will exhibit regular peaks at multiples values of the period (here 190 nm, 380 nm, 760 nm etc). The position of the first peak of the autocorrelation curve can be used to retrieve the average spacing (s), and its height estimates how marked the periodicity of the profile is. Axons were manually traced in ImageJ using polyline ROIs on 16 nm/pixel reconstructed images. The normalized autocorrelation curve of the corresponding intensity profile was calculated and plotted. The ImageJ script for generating autocorrelation curves from polyline ROIs is available at the following web address: https://github.com/cle-terrier/Process_Profiles. Autocorrelation curves of all tracings for a given labeling and condition were then averaged. The first non-zero peak of the averaged autocorrelation curve was fitted in Prism (Graphpad software) to estimate its position, providing the spacing value (s) and the corresponding error. The autocorrelation amplitude (height of the first peak) was estimated from the difference between the autocorrelation values at 192 nm (approximate position of the first peak) and 96 nm (approximate position of the first valley).

### EM of platinum replicas

Replicas on EM grids were mounted in a eucentric side-entry goniometer stage of a transmission electron microscope operated at 80 kV (model CM120; Philips) and images were recorded with a Morada digital camera (Olympus). Images were processed in Photoshop (Adobe) to adjust brightness and contrast and presented in inverted contrast. Tomograms were made by collecting images at the tilt angles up to ±25° relative to the plane of the sample with 5° increments. Images were aligned by layering them on top of each other in Photoshop. Measurement of actin braids spacing, length and width were performed on high magnification PREM views using ImageJ. Five lines were traced between consecutive braids (for braid spacing) or across a single braid, before and after eventual splitting (for braid width) and averaged to obtain each distance data point.

### Correlative SMLM/PREM

For correlative SMLM/PREM, unroofed and stained samples were first imaged by epifluorescence, then a STORM image was acquired (see above). At the end of the acquisition a objective-style diamond scriber (Leica) was used to engrave a ∼1 mm circle around the imaged area, and a low-magnification, 10X phase image was taken as a reference for relocation. After replica generation and prior to its release, the coated surface of each coverslip was scratched with a needle to make an EM-grid sized region of interest containing the engraved circle. Grids were imaged with low magnification EM to relocate the region that previously imaged by epifluorescence and SMLM, and high-magnification EM views were taken from the corresponding axonal region. A high-resolution SMLM reconstruction was mapped and aligned by affine transformation to the corresponding high-magnification EM view using the eC-CLEM plugin in ICY software (Paul-Gilloteaux et al., 2017).

### Data representation and statistical analysis

Intensity profiles, graphs and statistical analyses were generated using Prism. On bar graphs, dots (if present) are individual measurements, bars or horizontal lines represent the mean, and vertical lines are the SEM unless otherwise specified. Significances were tested using two-tailed unpaired non-parametric t-tests (two conditions) or one-way non-parametric ANOVA followed by Tukey post-test (3 or more conditions). In all figures, significance is coded as: ns non-significant, * p < 0.05, ** p < 0.01, *** p < 0.001. The number of data points and experiments for quantitative results is presented in Table S1 and S2. The measurements presented in Fig. 1 are from the same pooled dataset as the control condition in Fig. 3. Statistics from quantitative analyses are summarized in Supplementary Tables 1-3.

## Acknowledgments

We thank Matthew Rasband and Vann Bennett for antibody gifts; Fanny Boroni-Rueda for help with neuronal cultures; the NCIS imaging facility and Nikon Instruments for SMLM equipment; the IBPS cryo-EM facility for EM equipment; Subhojit Roy, Jeanne Lainé, Marc Bitoun, Ricardo Henriques, Manuel Théry, Laurent Blanchoin, Emmanuel Nivet and the NeuroCyto team members for discussions and critical reading of the manuscript. This work has been funded by the CNRS ATIP-AVENIR program (grant ATIP AO 2016 to CL); by Sorbonne Université, INSERM, Association Institut de Myologie core funding and the Agence Nationale de la Recherche (grant ANR14 CE12 0001 01 to SV).

## Author contributions

SV: Conceptualization, formal analysis, funding acquisition, investigation, methodology, visualization, writing – review and editing. SG, AJ and GC: formal analysis, investigation. CL: Conceptualization, formal analysis, funding acquisition, investigation, methodology, visualization, software, supervision, writing – original draft, review and editing.

## Author Information

Authors declare no competing interests. Correspondence and request for materials should be addressed to.

## Supplemental Information

**Supplementary Figure 1.**
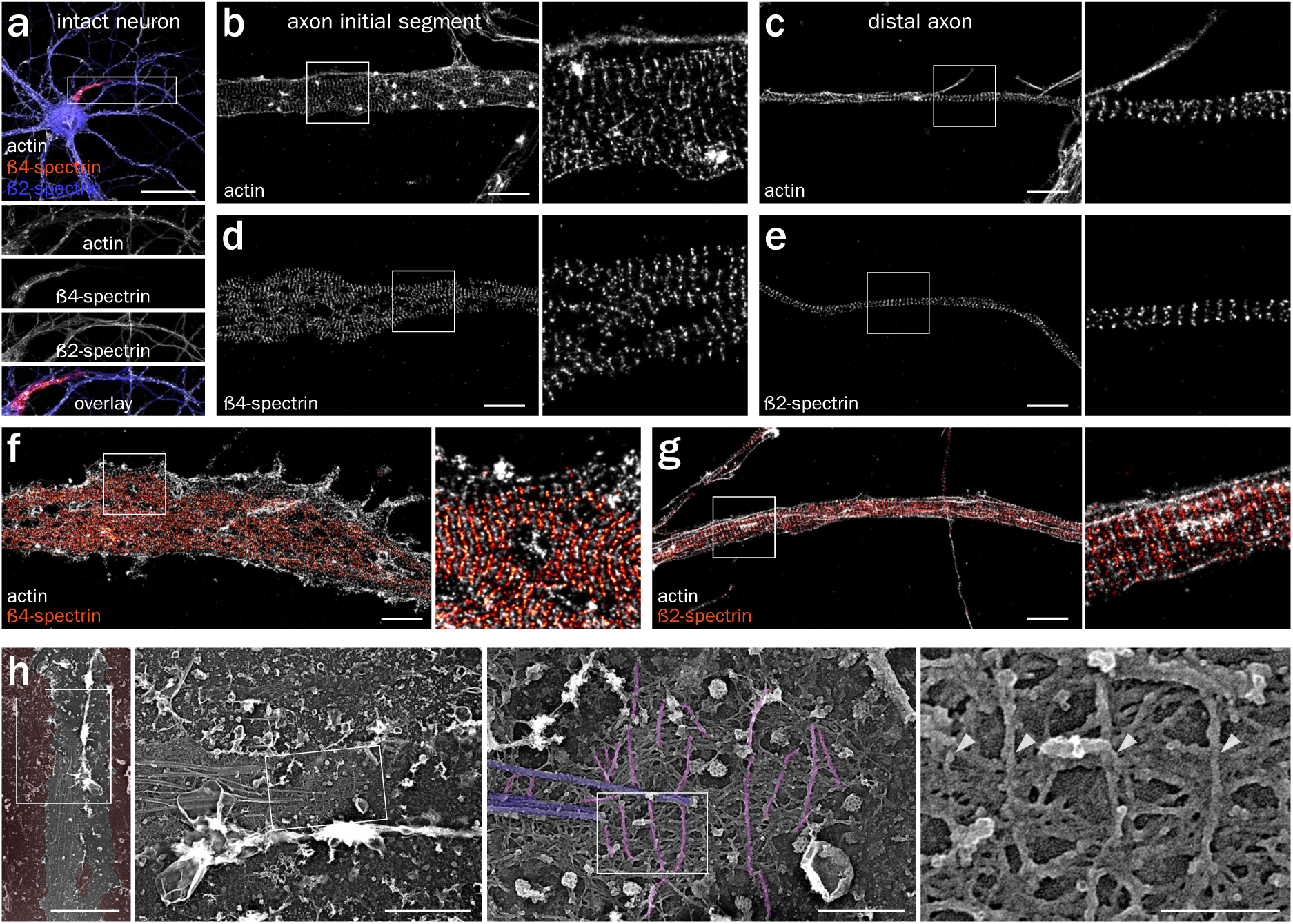
SMLM of the MPS along intact axons and additional PREM views of an unroofed axon. **(a)** Epifluorescence image of a neuron labeled for actin (gray), ß4-spectrin (orange) and ß2-spectrin (blue). **(b-c)** SMLM images showing actin rings along the AIS (b) and the distal axon (c). **(d-e)** SMLM images showing the periodic pattern of ß4-spectrin along the AIS (d) or ß2-spectrin along the distal axon (e). **(f-g)** SMLM images showing the alternate periodic pattern of actin (gray) and ß4-spectrin along the AIS (f, orange) or ß2-spectrin along the distal axon (g, orange). **(h)** PREM view of an unroofed axon showing the regularly spaced braids (magenta, arrowheads). Scale bars 40 μm (a), 2 μm (b-g), 5, 2, 0.5 and 0.2 μm (h, left to right).

**Supplementary Figure 2.**
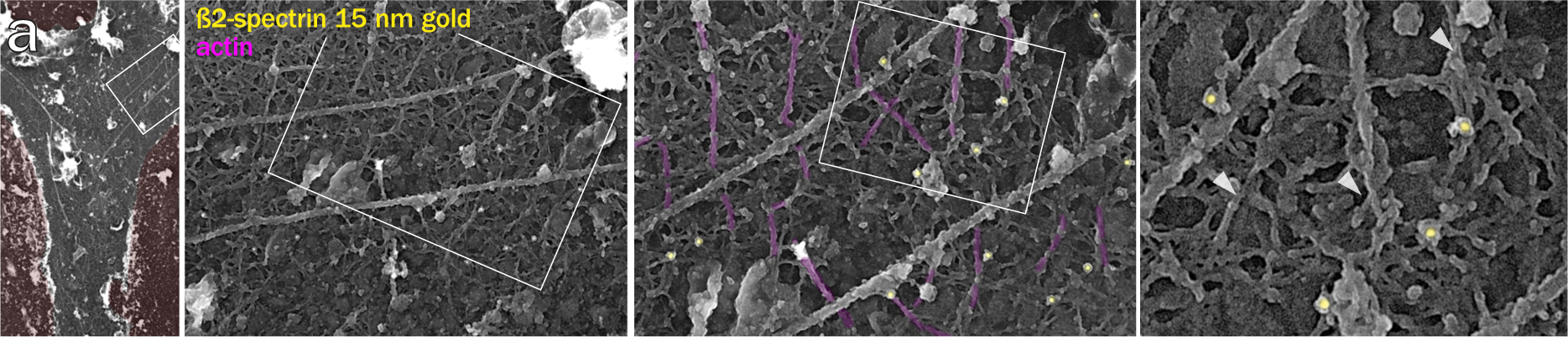
Immunogold labeling of ß2-spectrin. **(a)** PREM views of an unroofed axon immunogold-labeled (15 nm gold beads are pseudo-colored yellow) for ß2-spectrin between actin braids (magenta, arrowheads). Scale bars 2 µm, 1, 0.5, 0.2 µm (left to right).

**Supplementary Figure 3.**
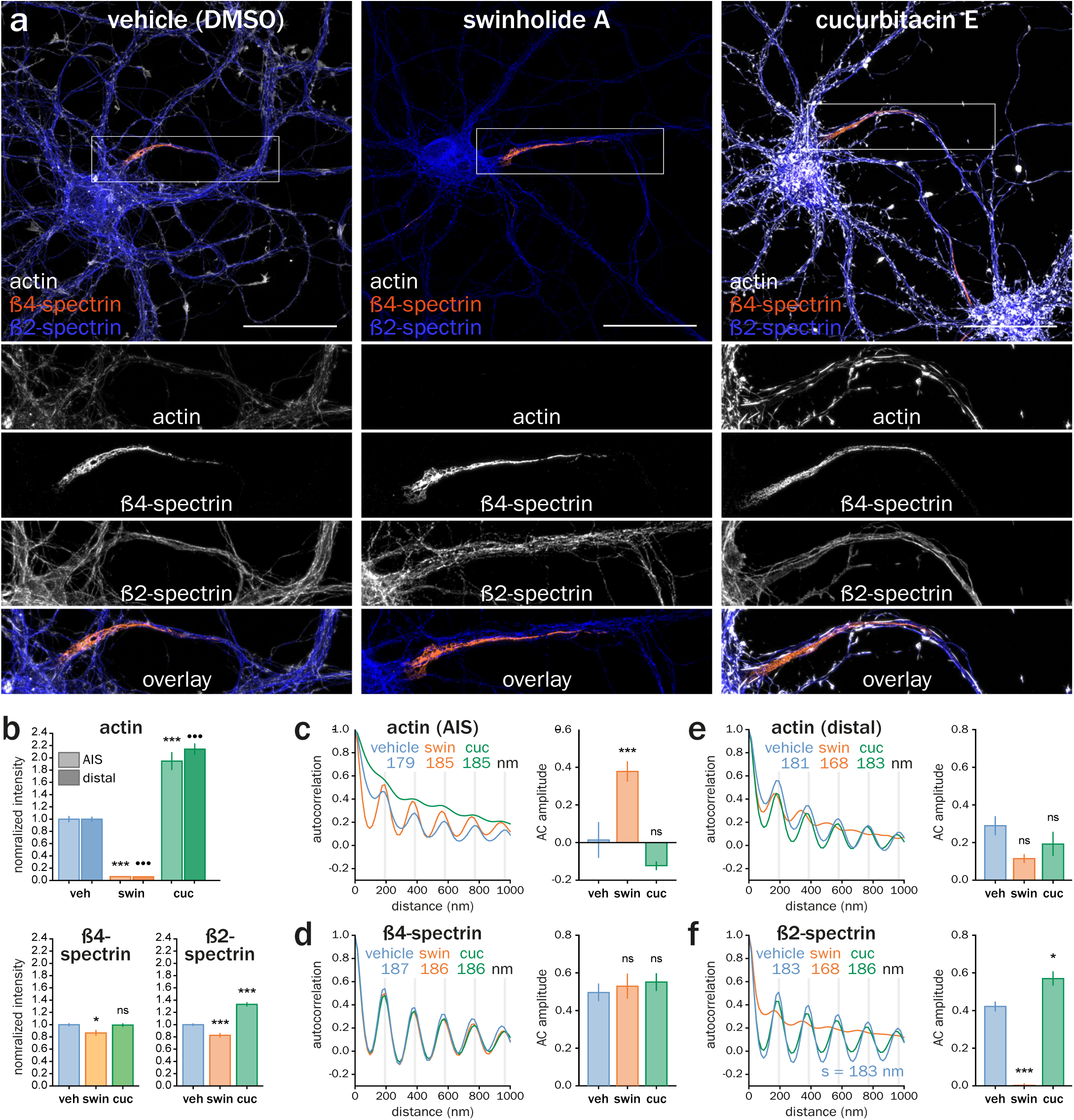
Epifluorescence images and quantification of drugs effects on MPS components. **(a)** Epifluorescence image of neurons treated for 3h with vehicle (DMSO 0.1%), swinholide A (100 nM) or cucurbitacin E (5 nM) and labeled for actin (gray), ß4-spectrin (orange) and ß2-spectrin (blue). **(b)** Top, normalized axonal labeling intensity for actin along the AIS (light) and distal axons (dark) measured on epifluorescence images of treated neurons. Bottom, same quantification for the ß4-and ß2-spectrin labeling. **(c-f)** Left, autocorrelation curve of the labeling after drug treatment for actin along the AIS (c), ß4-spectrin (d), actin along distal axons (e), or ß2-spectrin (f). Spacing measured from the autocorrelation curve is indicated. Right, measurement of the corresponding autocorrelation amplitude for the different treatments. Scale bar, 40 µm (a).

**Supplementary Figure 4.**
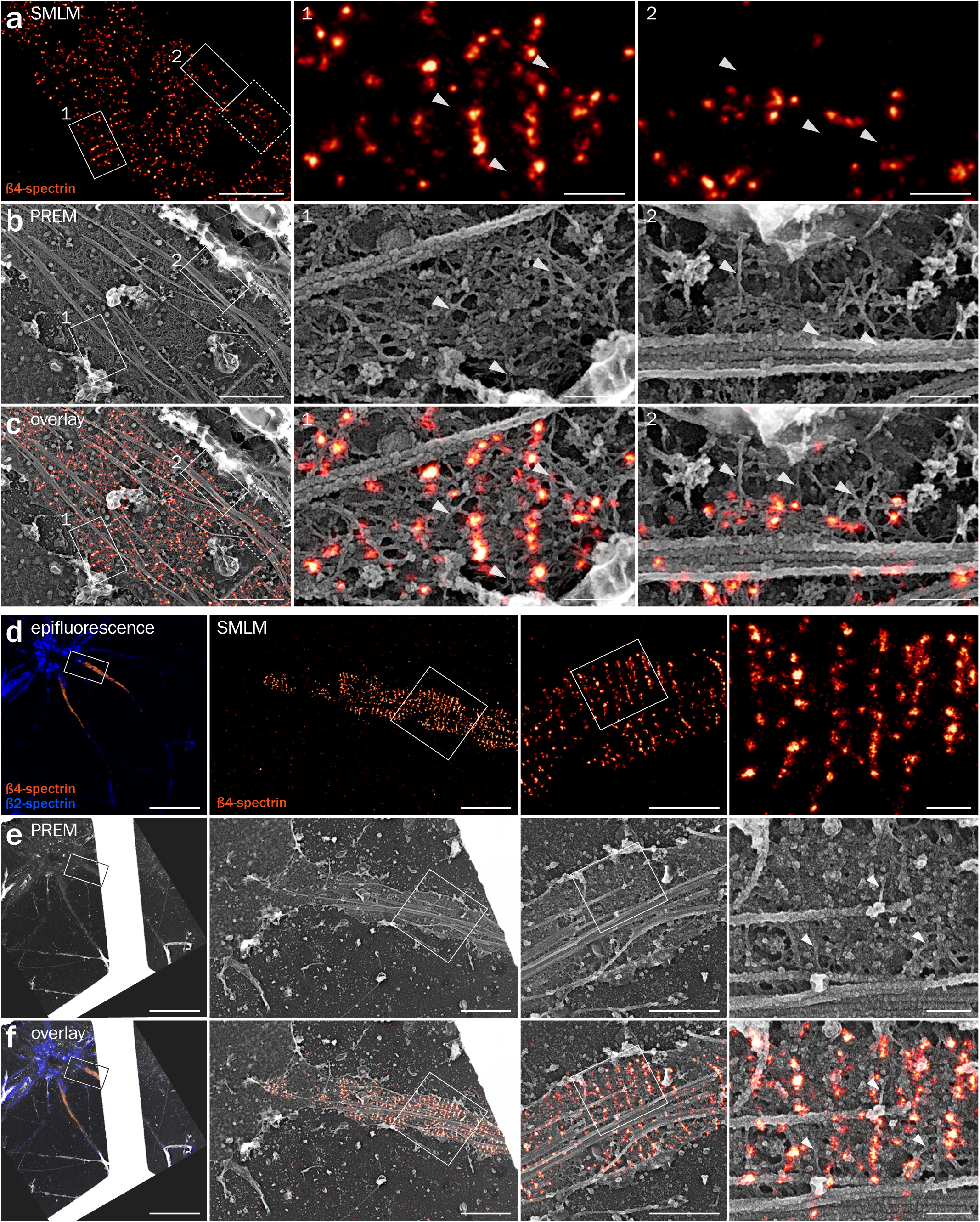
Additional examples for correlative SMLM/PREM of ß4-spectrin. **(a-c)** Additional zooms from the SMLM/PREM image of ß4-spectrin along the axon shown in Fig. 5. Zoom shown in Fig. 5 is highlighted by a dotted box on the image on the left, and additional zooms 1-2 presented on the right are highlighted by solid boxes. SMLM image of ß4-spectrin labeling (a), corresponding PREM view (b) and overlay of the SMLM and PREM images (c). Scale bars 1, 0.2, 0.2 µm (from left to right). **(d)** Left, epifluorescence image of another unroofed neuron labeled for ß4-spectrin (orange) and ß2-spectrin (blue). Right, SMLM images of the unroofed axon labeled for ß4-spectrin. **(e)** Corresponding PREM views of the same unroofed neuron and axon (actin braids, arrowheads). **(f)** Overlay of the SMLM image and PREM views. Scale bars 20, 2, 1, 0.2 µm (from left to right).

**Supplementary Figure 5.**
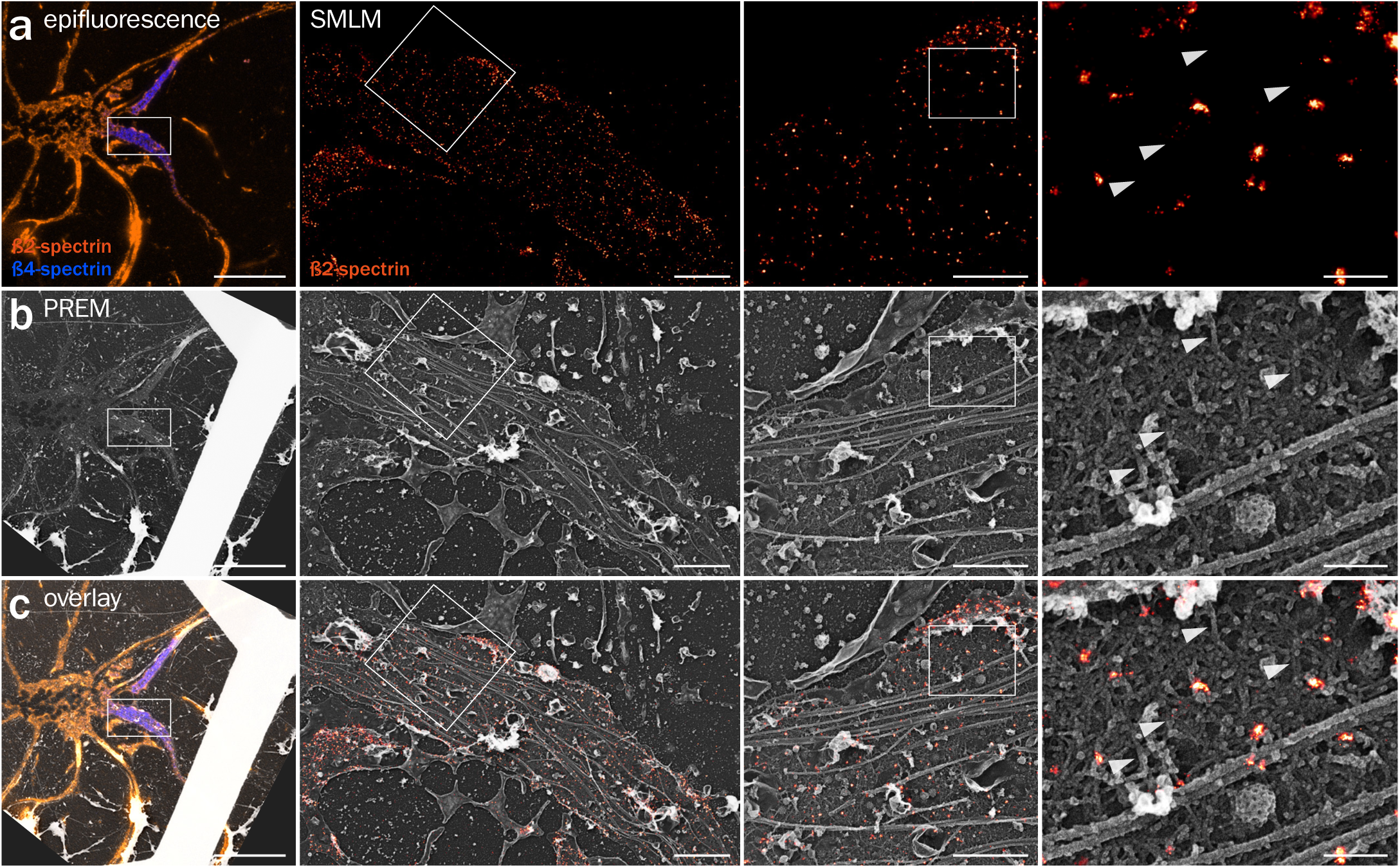
Correlative SMLM/PREM of ß2-spectrin. **(a)** Left, epifluorescence image of an unroofed neuron labeled for ß2-spectrin (orange) and ß4-spectrin (blue). Right, SMLM images of the un-roofed axon labeled for ß2-spectrin. **(b)** Corresponding PREM views of the same unroofed neuron and axon (actin braids, arrowheads). **(c)** Overlay of the SMLM image and PREM views. Scale bars 20, 2, 1, 0.2 µm (from left to right).

**Supplementary Figure 6.**
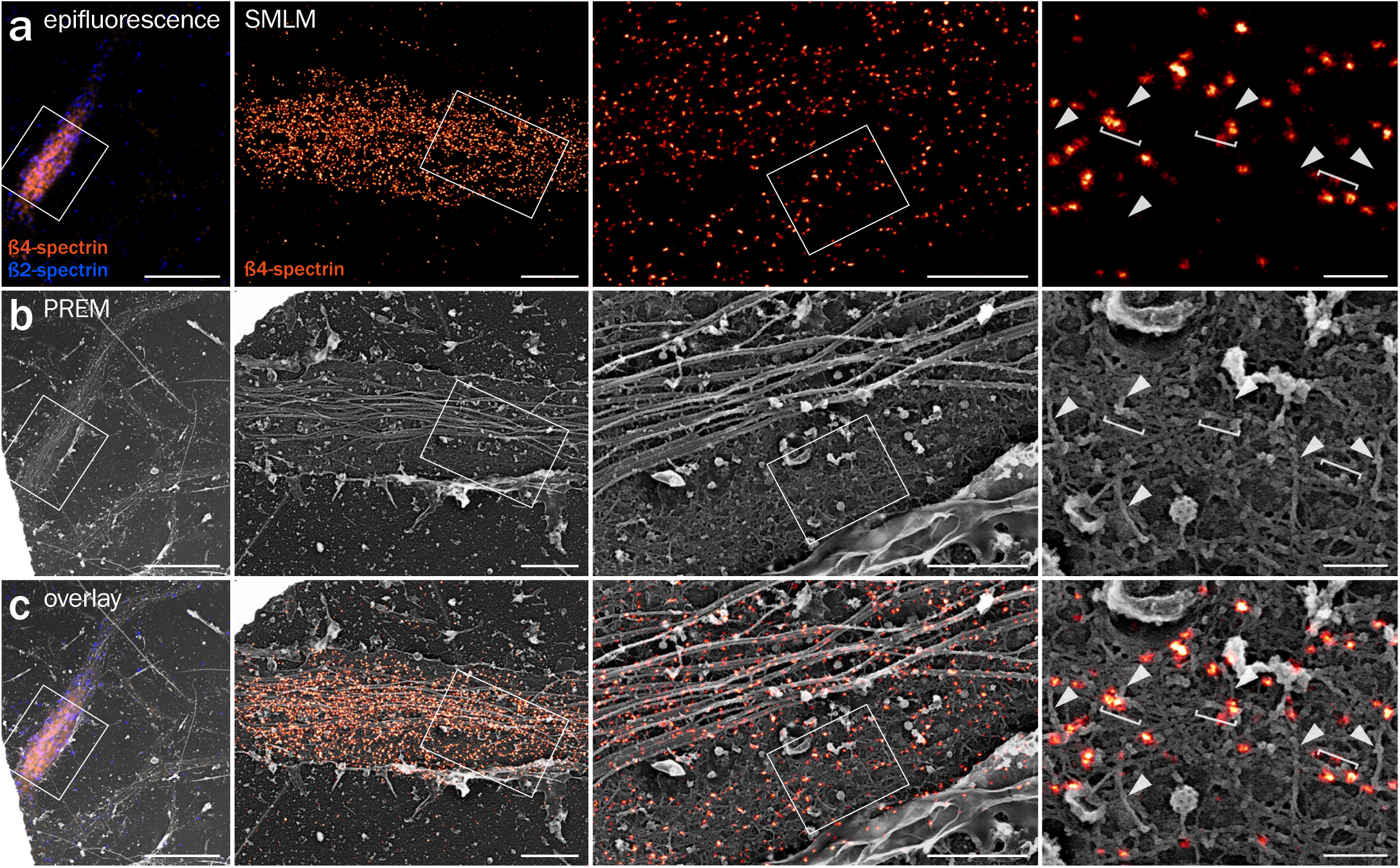
Correlative SMLM/PREM of pMLC. **(a)** Left, epifluorescence image of an unroofed neuron labeled for pMLC (orange) and ankyrin G (blue). Right, SMLM images of the unroofed proximal axon labeled for pMLC. **(b)** Corresponding PREM views of the same unroofed neuron and axon. **(c)** Overlay of the SMLM image and PREM views showing pMLC apposed to actin breads (arrowheads). Scale bars 20, 2, 1, 0.2 µm (from left to right).

## Supplementary Tables

**Supplementary Table 1.** Data statistics summary for fluorescence microscopy quantifications.

| Labeling | Treatment | Intensity along axons from epifluorescence |  |  |  |  | Autocorrelation of axonal segments from SMLM |  |  |  |  |  |  |
| --- | --- | --- | --- | --- | --- | --- | --- | --- | --- | --- | --- | --- | --- |
|  |  | Normalized intensity |  | Number of data points |  |  | Spacing s (μm) |  | Amplitude |  | Number of data points |  |  |
|  |  | Mean | SEM | Tracings | Images | Indep. exp | Fit value | Error | Mean | SEM | Tracings | Images | Indep. exp |
| actin (AIS) | vehicle | 1,000 | 0,051 | 77 | 34 | 5 | 0,179 | 0,007 | 0,014 | 0,095 | 18 | 14 | 4 |
|  | unroofed |  |  |  |  |  | 0,187 | 0,002 | 0,302 | 0,043 | 42 | 32 | 6 |
|  | swin | 0,061 | 0,006 | 46 | 30 | 3 | 0,185 | 0,003 | 0,378 | 0,054 | 20 | 9 | 3 |
|  | cuc | 1,948 | 0,146 | 65 | 24 | 3 | 0,142 | 0,287 | -0,123 | 0,023 | 8 | 9 | 3 |
| actin (distal) | ctrl | 1,000 | 0,042 | 451 | 34 | 5 | 0,181 | 0,003 | 0,289 | 0,051 | 52 | 13 | 4 |
|  | swin | 0,061 | 0,006 | 177 | 30 | 3 | 0,168 | 0,012 | 0,115 | 0,023 | 15 | 9 | 3 |
|  | cuc | 2,142 | 0,095 | 283 | 24 | 3 | 0,183 | 0,004 | 0,192 | 0,064 | 27 | 9 | 3 |
| β4-spectrin | ctrl | 1,000 | 0,026 | 107 | 56 | 6 | 0,187 | 0,002 | 0,497 | 0,047 | 43 | 14 | 3 |
|  | unroofed |  |  |  |  |  | 0,191 | 0,002 | 0,527 | 0,070 | 16 | 9 | 2 |
|  | swin | 0,867 | 0,047 | 74 | 52 | 4 | 0,186 | 0,002 | 0,530 | 0,067 | 25 | 12 | 2 |
|  | cuc | 0,993 | 0,029 | 105 | 46 | 5 | 0,186 | 0,002 | 0,552 | 0,047 | 33 | 14 | 3 |
| β2-spectrin | ctrl | 1,000 | 0,019 | 777 | 66 | 7 | 0,184 | 0,001 | 0,422 | 0,026 | 60 | 18 | 4 |
|  | unroofed |  |  |  |  |  | 0,185 | 0,003 | 0,367 | 0,041 | 30 | 15 | 2 |
|  | swin | 0,677 | 0,027 | 450 | 62 | 5 | 0,163 | 0,017 | 0,003 | 0,010 | 29 | 15 | 3 |
|  | cuc | 1,285 | 0,031 | 502 | 56 | 5 | 0,188 | 0,001 | 0,570 | 0,038 | 31 | 11 | 2 |

**Supplementary Table 2.**
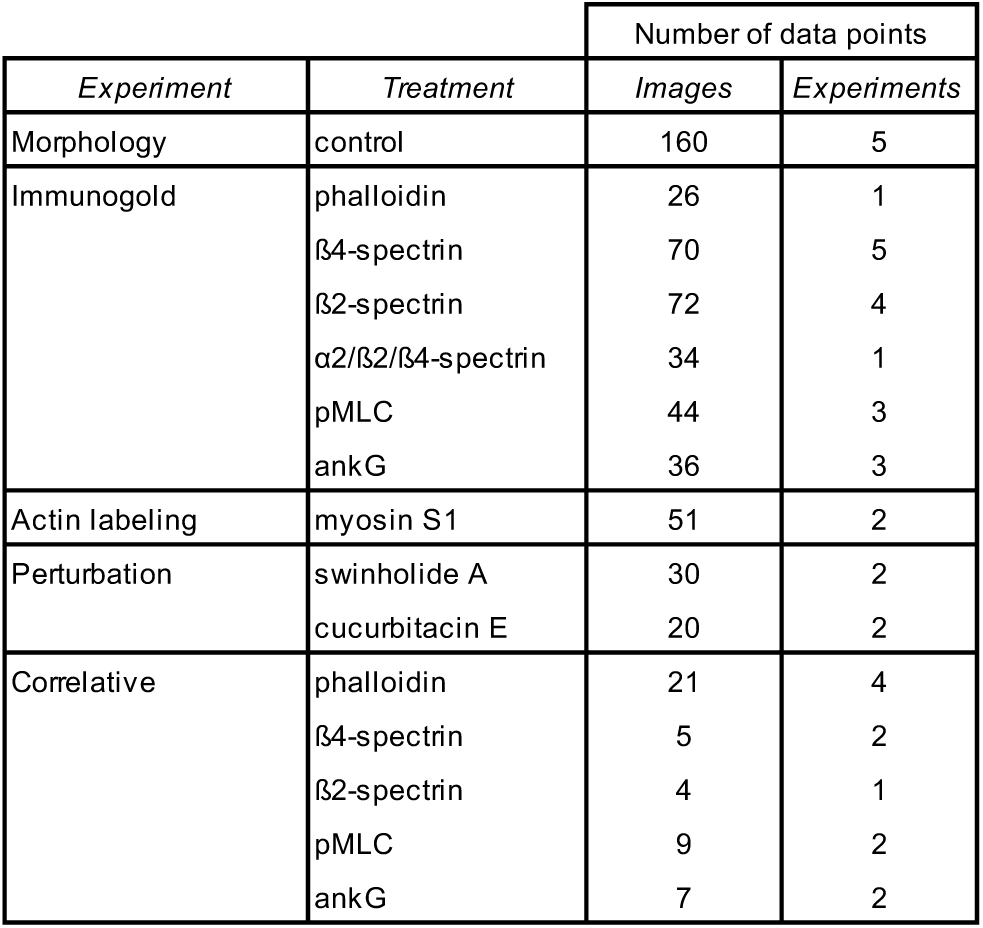
Summary of image numbers for electron microscopy.

**Supplementary Table 3.** Data statistics summary for electron microscopy quantifications.

| Measurement | Treatment | Analysis from PREM views |  |  |  |  |
| --- | --- | --- | --- | --- | --- | --- |
|  |  | Value (nm) |  | Number of data points |  |  |
|  |  | Mean | SEM | Filaments | Images | Exp |
| filaments spacing | control | 183,8 | 4,5 | 50 | 10 | 5 |
| filaments length | control | 688,6 | 41,2 | 45 | 6 | 5 |
|  | myosin S1 | 1130,0 | 44,5 | 76 | 13 | 2 |
| filaments thickness | braid | 18,5 | 0,2 | 90 | 31 | 8 |
|  | split braid | 10,2 | 0,3 | 52 | 18 | 8 |
|  | dendrite | 9,9 | 0,2 | 60 | 5 | 3 |
|  | microtubule | 31,0 | 0,7 | 33 | 6 | 5 |

## Captions for Supplementary Movies 1 to 7

**Supplementary Movie 1. Tomogram corresponding to Fig. 1j.**

PREM tomogram of an unroofed axon. Actin braids appear in magenta during the movie, then the spectrin mesh in yellow, and microtubules in blue.

**Supplementary Movie 2. Tomogram corresponding to Fig. 1k.**

PREM tomogram of an unroofed axon. Actin braids appear in magenta during the movie, then the spectrin mesh in yellow, and microtubules in blue.

**Supplementary Movie 3. Tomogram corresponding to Supplementary Figure 1h.**

PREM tomogram of an unroofed axon. Actin braids appear in magenta during the movie, then the spectrin mesh in yellow, and microtubules in blue.

**Supplementary Movie 4. Tomogram corresponding to Fig. 2g.**

PREM tomogram of an unroofed axon. Actin braids appear in magenta during the movie, then the spectrin mesh in yellow, and microtubules in blue.

**Supplementary Movie 5. Correlative PREM/SMLM for actin corresponding to Fig 5a-c.**

The unroofed neuron is shown with successive epifluorescence image (ß2-spectrin in green, ß4-spectrin in red, actin in blue), low-magnification PREM image (grid appears white), SMLM image (actin in orange) and high-magnification PREM image of the proximal axon. The high-magnification PREM image of the axon is then superimposed with the SMLM image (actin in orange).

**Supplementary Movie 6. Correlative PREM/SMLM for ß4-spectrin corresponding to Supplementary Figure 4d-f.**

The high-magnification PREM image of the axon is superimposed with the SMLM image (ß4-spectrin in orange).

**Supplementary Movie 7. Correlative PREM/SMLM for ß2-spectrin corresponding to Supplementary Figure 5a-c.**

The high-magnification PREM image of the axon is superimposed with the SMLM image (ß2-spectrin in orange).

